# Hebbian priming of human spinal motor learning

**DOI:** 10.1101/2023.02.17.528541

**Authors:** Jonas Rud Bjørndal, Mikkel Malling Beck, Lasse Jespersen, Lasse Christiansen, Jesper Lundbye-Jensen

**Affiliations:** Movement & Neuroscience, Department of Nutrition, Exercise and Sports (NEXS), University of Copenhagen, Nørre Allé 51, 2200 Copenhagen N, Denmark; Danish Research Centre for Magnetic Resonance, Centre for Functional and Diagnostic Imaging and Research, Copenhagen University Hospital Amager and Hvidovre, Kettegård Allé 30, 2650 Hvidovre, Denmark

**Keywords:** Plasticity/neuroplasticity/Hebbian plasticity/Hebbian Priming, motor learning, paired stimulation, Transcranial magnetic stimulation

## Abstract

Learning or relearning of motor skills requires plasticity in relevant neural circuits. Motor recovery following lesions to the corticospinal system can be augmented through neuromodulation techniques targeting the affected or compensatory neural circuits. By repeatedly pairing transcranial magnetic stimulation of the primary motor cortex (M1) and motoneuronal electrical stimulation (i.e., paired corticomotoneuronal stimulation, PCMS) timed to arrive at the corticomotoneuronal (CM) synapses in close temporal proximity, spike-timing-dependent bidirectional changes in CM transmission can be induced in humans (Taylor & Martin, 2009). PCMS-induced increases in CM transmission have been demonstrated to transiently improve motor control in patients with spinal cord injury (Bunday & Perez 2012), whereas effects on the malleability of neural circuits are entirely unexplored. We hypothesized that PCMS can prime mechanisms of subsequent motor learning exclusively when directed to the neural circuitry underpinning the motor behavior. In three experiments, we provide the first evidence (‘Experiment I’) and a double-blinded, sham-controlled replication (‘Experiment II’) that PCMS targeting the spinal CM synapses can prime subsequent learning of rapid finger movements relying on spinal neuroplasticity. Finally, we demonstrate that the effects of PCMS are circuit-specific and bidirectional. When PCMS was timed to arrive at a facilitatory interval in M1 but an inhibitory interval at the CM synapses subsequent learning was transiently impeded (‘Experiment III’). Taken together, our results provide proof-of-principle that non-invasively induced plasticity governed by Hebbian learning rules interacts with experience-dependent plasticity in the spinal cord with positive implications for motor learning. Our results offer a mechanistic rationale for priming sensorimotor training with individualized PCMS to enhance the effects of motor practice in neurorehabilitation.

**Highlights:**

- Paired corticomotoneuronal stimulations (PCMS) promote ballistic motor learning and facilitate corticospinal excitability compared to rest and sham protocols.
- A double-blinded sham experiment replicates priming effects of PCMS on ballistic motor learning and demonstrates long-term benefits of combined PCMS and motor practice.
- The facilitating effect of PCMS on ballistic motor learning is circuit-specific with superior effects on ballistic motor learning after facilitating PCMS compared to control protocols.

## Results & Discussion

Across organisms, motor learning is governed by experience-dependent plasticity in relevant neural circuits along the neuroaxis dependent on the task demands (Shmuelof & Krakauer, 2011). These intrinsic processes of learning may be modulated extrinsically by non-invasive neuromodulation techniques. In a series of three experiments, we used non-invasive brain and peripheral nerve stimulation to prime subsequent spinal motor learning in humans. We found that motor learning involving ballistic index finger movements, a motor skill contingent on spinal plasticity, was improved by preceding paired corticomotoneuronal stimulation (PCMS), a well-established human model for exogenous induction of spike-timing-dependent plasticity (STDP) (Experiment I). We confirmed the findings in a subsequent double-blinded experiment and expanded the clinical relevance by demonstrating that performance remained facilitated seven days after initial priming (Experiment II).

Finally, we demonstrated that effects of PCMS on ballistic learning were bidirectional and circuit-specific in that only paired stimulations timed to facilitate corticomotoneuronal transmission increased learning. In contrast, paired stimulation timed to inhibit corticomotoneuronal synaptic transmission delayed acquisition (Experiment III).

### Experiment 1: Hebbian priming of ballistic motor learning

In Experiment I (N=26), we individualized PCMS so that the descending corticospinal volleys elicited by TMS of the hand area of the primary motor cortex (M1) arrived at the corticomotoneuronal pre-synapse 2 ms before a peripherally triggered antidromic volley in the motor axons arrived at the post-synapse (referred to as the ‘PCMS+’)(Figure 1A, and see Supplementary information, Figure S1 for details). This protocol has consistently been shown to increase corticospinal excitability in humans (Bunday & Perez, 2012; Shulga et al., 2015; Taylor & Martin, 2009). We found that priming ballistic motor learning with 100 paired stimuli resulted in superior learning evidenced by better ballistic performance at the end of practice compared to controls who only performed motor practice and received no paired stimuli (‘Rest’) (Figure 1B). This was supported statistically by linear mixed effects model (LMM) showing a significant GROUP x TIME interaction (F_(3,4038)_=52, p<0.001) on peak index finger acceleration. We found that PCMS led to a significantly larger improvement in performance from the first practice block (B1) to the last practice block (B3) (i.e. online learning, PCMS+: +22.28%±1.0 vs. Rest: 10.93%±1.0, p<0.001, Figure 1C). Performance did not differ between groups at baseline (p=0.727), suggesting that the differences that emerged during practice were not due to general differences in task proficiency but likely could be ascribed to PCMS+.

**Figure 1.**
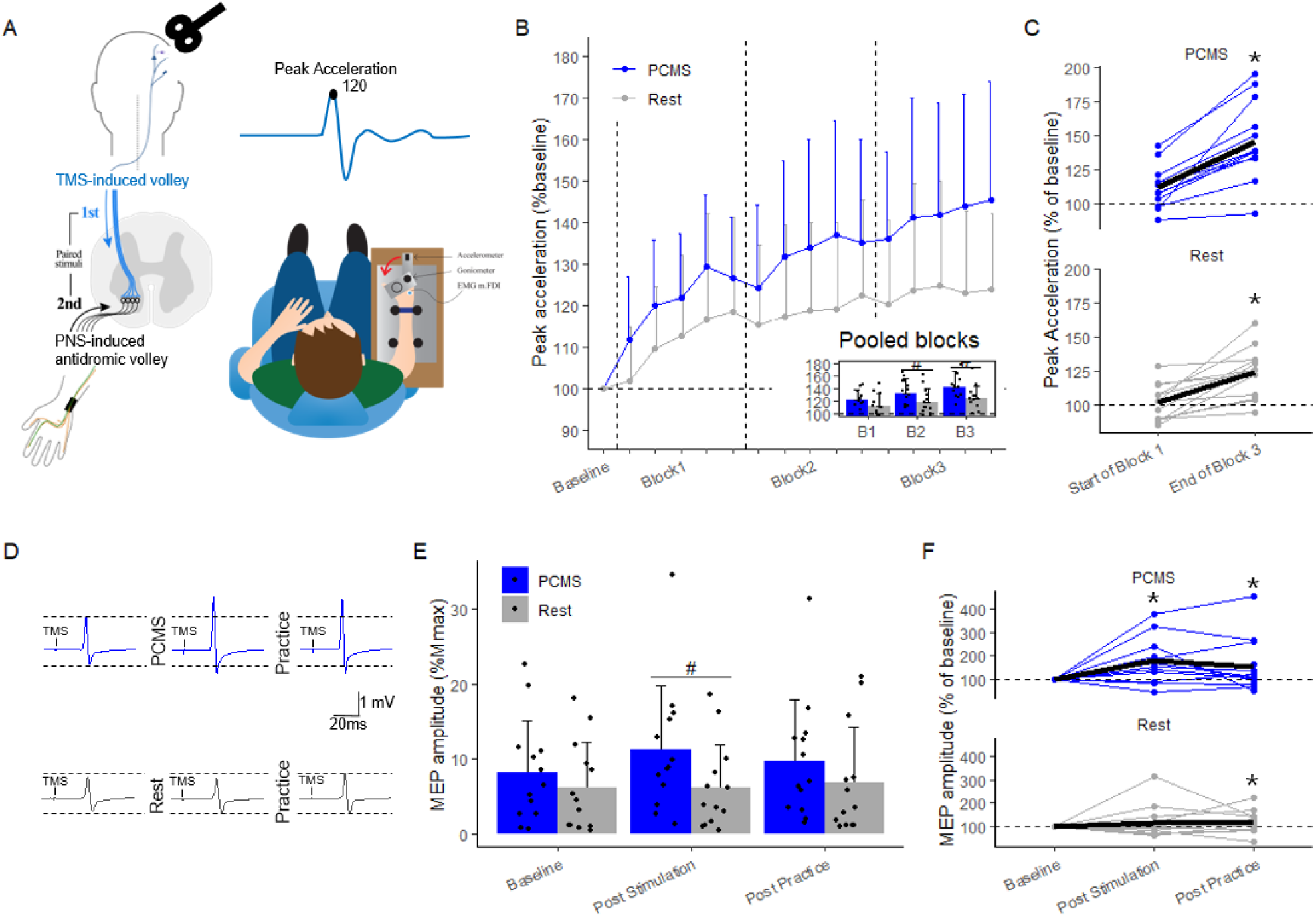
Experiment I: Effects of PCMS and rest on ballistic motor learning and corticospinal excitability. **A)** Between-group design, 26 participants were randomized to two groups, PCMS (blue) or Rest (red). **B)** Learning curves after PCMS or Rest conditions, each datapoint represents the mean of 10 trials of the ballistic index finger flexion task relative to individual mean at baseline. The inserted bar chart shows group means across entire practice blocks with individual dots for each participant. **C)** Individual data from the two conditions showing the mean of the 10 first trials of practice, compared to the mean of the last 10 trials of practice. The black line represents the mean performance across participants. **D)** Example of single subject data, raw MEP traces after PCMS or Rest. **E)** Motor evoked potentials in percentage of M_max_ amplitude were used to quantify corticospinal excitability. **F)** Acute individual response (MEP % baseline) to PCMS or sham conditions. The black line represents the mean MEP amplitudes across participants. #p<0.05 indicates a significant between-group comparison. *p<0.05 indicates significant within-group comparisons.

For both groups, we also probed corticospinal excitability through motor evoked potentials (MEP) elicited by TMS of the M1 and recorded from the first dorsal interosseous (FDI) muscle at baseline, post stimulation and post practice (Figure 1D). We observed no baseline difference in terms of MEP amplitude expressed as the ratio to M_max_ amplitude (PCMS+= 0.08±0.06 vs. ‘Rest’ 0.06±0.05, p=0.26) (Figure 1E). In line with previous work, PCMS+ increased MEP amplitudes as compared to the resting condition (Figure 1E) evident as a significant GROUPxTIME interaction (F_(2,1766)_=10.9, p<0.001). Post-hoc comparisons revealed that PCMS+ induced larger increases in MEP amplitudes from baseline to post stimulation compared to ‘Rest’ (PCMS = 103.5%±11.3 vs. Rest =26.4%±12.4, p<0.01) and that there were no differences at baseline (p=0.23).

In Experiment I, we provide first evidence of an interaction between stimulation-induced and practice-dependent neuroplasticity leading to improved ballistic learning and coinciding increases in corticospinal excitability. Since previous results have shown that PCMS protocols do not increase maximal voluntary contraction (D’Amico et al., 2018; Dongés et al., 2019) the immediate effects of PCMS on motor performance may be limited to dexterous motor functions. We found a positive priming effect on ballistic motor *learning* measured as peak index finger acceleration in PCMS compared to rest. Without any directional movement constraints, peak acceleration in a specific direction can be increased by optimizing movement direction (likely a cortical phenomenon) as well as acceleration *per se* by improving fast, coordinated activation of agonist spinal motoneurons. In line with previous experiments, we confined ballistic index finger movements to one plane to emphasize the spinal contribution to learning (Giesebrecht et al., 2012). Consequently, the observed increase in peak acceleration can be assumed to largely reflect improved efficacy of direct activation of spinal motoneurons.

In primates, the motoneurons innervating hand muscles are to a large extent excited through corticospinal projections with monosynaptic connections from the cortical hand area of the precentral gyrus (Sinopoulou et al., 2022). The direct corticomotoneuronal projections are also the prime candidate signaling pathway for TMS-evoked excitation of motoneurons (see e.g. Siebner et al., 2022) and our results replicate previous findings that PCMS targeting the spinal cord increases the motor response to TMS over M1 (see Christiansen & Perez, 2018, for review).

We found short-term facilitatory effects of PCMS+ compared to rest on ballistic motor learning in able-bodied participants. However, the long-term effects of the PCMS-priming remained unexplored. Furthermore, we observed a numerical difference in average performance in the first block of motor practice after ‘Rest’ that we ascribe to a decrease in vigilance during the resting period. To ensure that the observed priming effect from PCMS+ on motor learning was not due to differences in state of alertness affecting motor performance, we introduced a sham protocol that mimicked the perceptual experience of PCMS+ in the following Experiment II. This double-blinded, sham-controlled experiment that included a 1-week retention test was performed to replicate our results from Experiment I and to assess the long-term motor effects caused by Hebbian priming.

### Experiment II: Double-blinded evidence of long-lasting behavioral benefits from priming PCMS

In twenty participants, we replicated the behavioral findings from Experiment I in a double-blinded, sham-controlled study. The PCMS protocol was similar to the one used in Experiment I whereas sham stimulation consisted of PNS delivered just above perceptual threshold and the TMS coil turned upside down. A LMM from Experiment II showed a significant interaction effect of GROUP and TIME (F_(4,3882)_=14.6, p<0.001). Post-hoc comparisons showed that there were no differences between PCMS+ and SHAM in peak acceleration at baseline (p=0.66), and that PCMS+ led to a significantly greater change in performance from B1 to B3 (i.e. online learning, PCMS+: +17.36%±0.8 vs. SHAM: 11.75%±0.8, p<0.001).

In addition, a 7-day follow-up test was included in Experiment II to assess long-term effects of PCMS compared to SHAM (Figure 2C) on retention following motor learning. Between-group comparison showed that the PCMS+ group performed significantly better than the SHAM group throughout 50 practice trials at day 7 (PCMS+: +120.58%±0.8 vs. SHAM: 115.56%±0.9, p<0.001).

**Figure 2.**
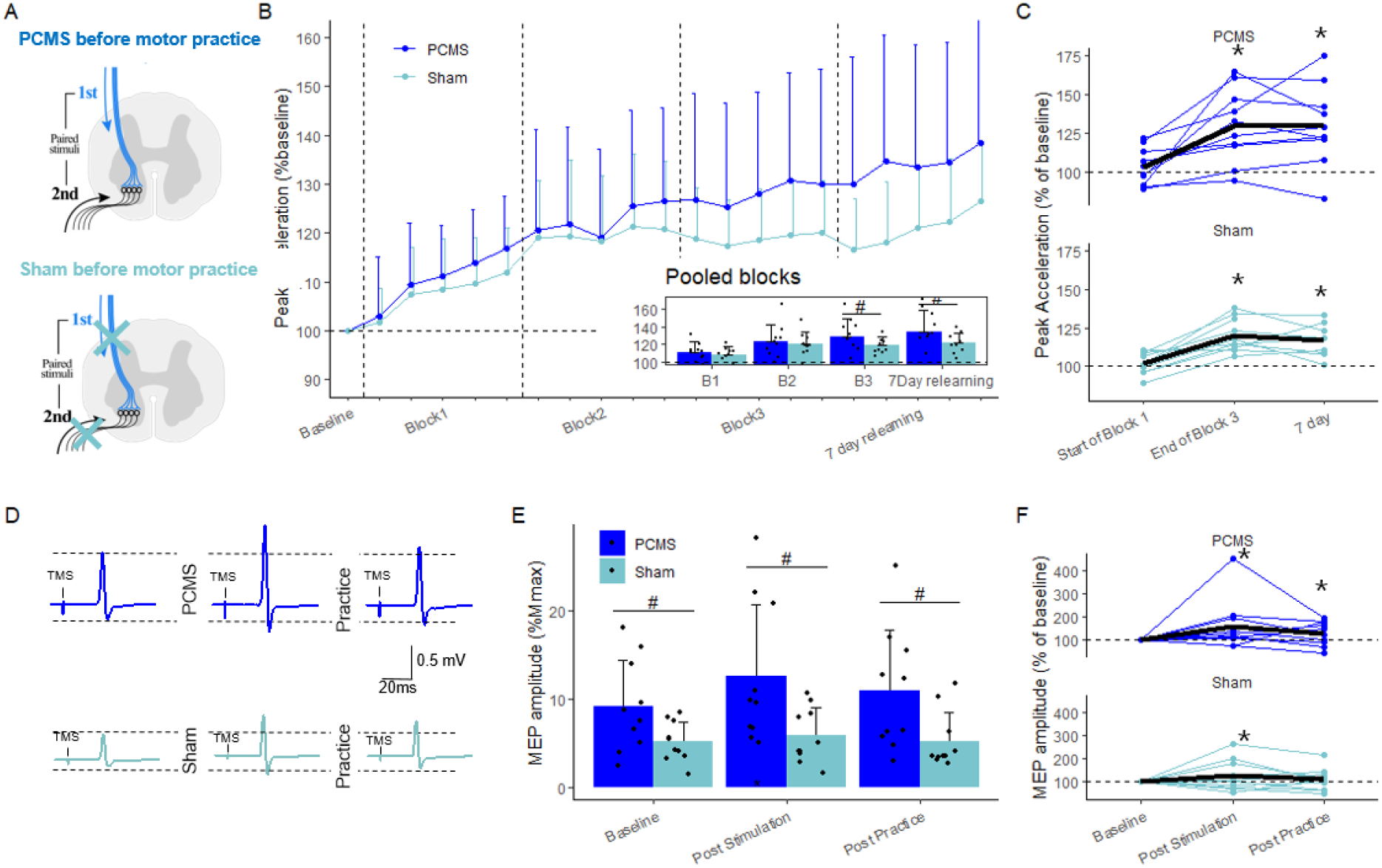
Experiment II: Effects of PCMS and sham protocols on ballistic motor learning and corticospinal excitability. **A)** Double-blinded, sham-controlled design, 20 participants were randomly assigned to one of two treatments, PCMS (blue) or Sham (light blue). **B)** Learning curves after PCMS or Sham conditions, each datapoint represents the mean of 10 trials relative to individual mean at baseline. The insert shows group means across entire practice blocks with individual dots for each participant. **C)** Individual data showing the mean of the first and last ten trials of practice along with the first ten trials seven days later. The black line represents the mean performance across participants. **D)** Example of single subject data, raw MEP traces after PCMS condition. **E)** Mean (bar) and individual (dots) amplitudes of motor evoked potentials expressed as the percentage of M_max_ amplitude. **F)** Individual average MEP amplitudes expressed relative to baseline after PCMS or Sham (top) and again following practice (bottom). The black line represents the mean MEP amplitudes across participants. #p<0.05 indicates significant between-group comparison. *p<0.05 indicates significant within-group comparison.

The assessment of corticospinal excitability revealed a pattern in part aligned with Experiment 1 and previous results (Bunday & Perez, 2012; Christiansen et al., 2018): LMM revealed a significant GROUPxTIME interaction (F_(2,1457)_=5.9, p<0.01). The two groups were not entirely matched in terms of MEP amplitude expressed as ratio to the M_max_, at baseline (PCMS = 0.09±0.05 vs. SHAM = 0.05±0.02, p=0.03) (Figure 2D-E). Despite of higher MEP amplitudes in PCMS compared to SHAM at baseline, post-hoc comparisons revealed that PCMS induced relatively larger increases in MEP amplitudes from baseline to post stimulation (PCMS = 58.2%±8.1 vs. SHAM =25.3%±8.1, p<0.01).

The results from Experiment II replicated the beneficial effects of Hebbian priming on ballistic motor learning. During sham, PNS was delivered just above perceptual threshold and the TMS coil was turned upside down. Thereby PNS did not elicit an antidromic volley and TMS did not elicit descending volleys while skin and scalp sensations along with an acoustic experience were maintained. Differences in the effects of PCMS+ and SHAM can consequently be assumed to reflect the intended pairing at the spinal level and not peripheral confounds (Figure 2B). In addition to replicating the acute effect on ballistic skill acquisition, we found that the benefits of PCMS persist after 7 days, i.e., a long-lasting effect on retention following combined PCMS and motor practice. This suggests that PCMS+ results in long lasting effects on motor learning that may have therapeutical relevance.

A third experiment was performed to investigate whether the effects of PCMS on motor learning were circuit specific (only when targeting CM synapses) and bidirectional (capable of both priming and inhibiting ballistic learning depending on spike-timing).

### Experiment III: Timing-dependent and network-specific effects of PCMS

Cellular STDP is characterized by bidirectional after-effects depending on both temporal proximity and order of pre- and postsynaptic spiking (Bi & Poo, 1998). To investigate if a similar learning rule applied to the network mediating the priming effects on ballistic motor learning shown in Experiment I and II, we conducted a within-subject cross-over experiment. In Experiment III, we compared the effects of PCMS+ to a protocol previously shown to induce inhibitory effects at the level of the CM synapse (‘PCMS-’, interarrival interval of 15ms, Taylor & Martin, 2009) and a coupled control protocol where the peripheral stimulation preceded the cortical stimulation with 100ms, i.e., well outside the window of spinal interactions (PCMS_coupled-control_) (Figure 3A).

**Figure 3.**
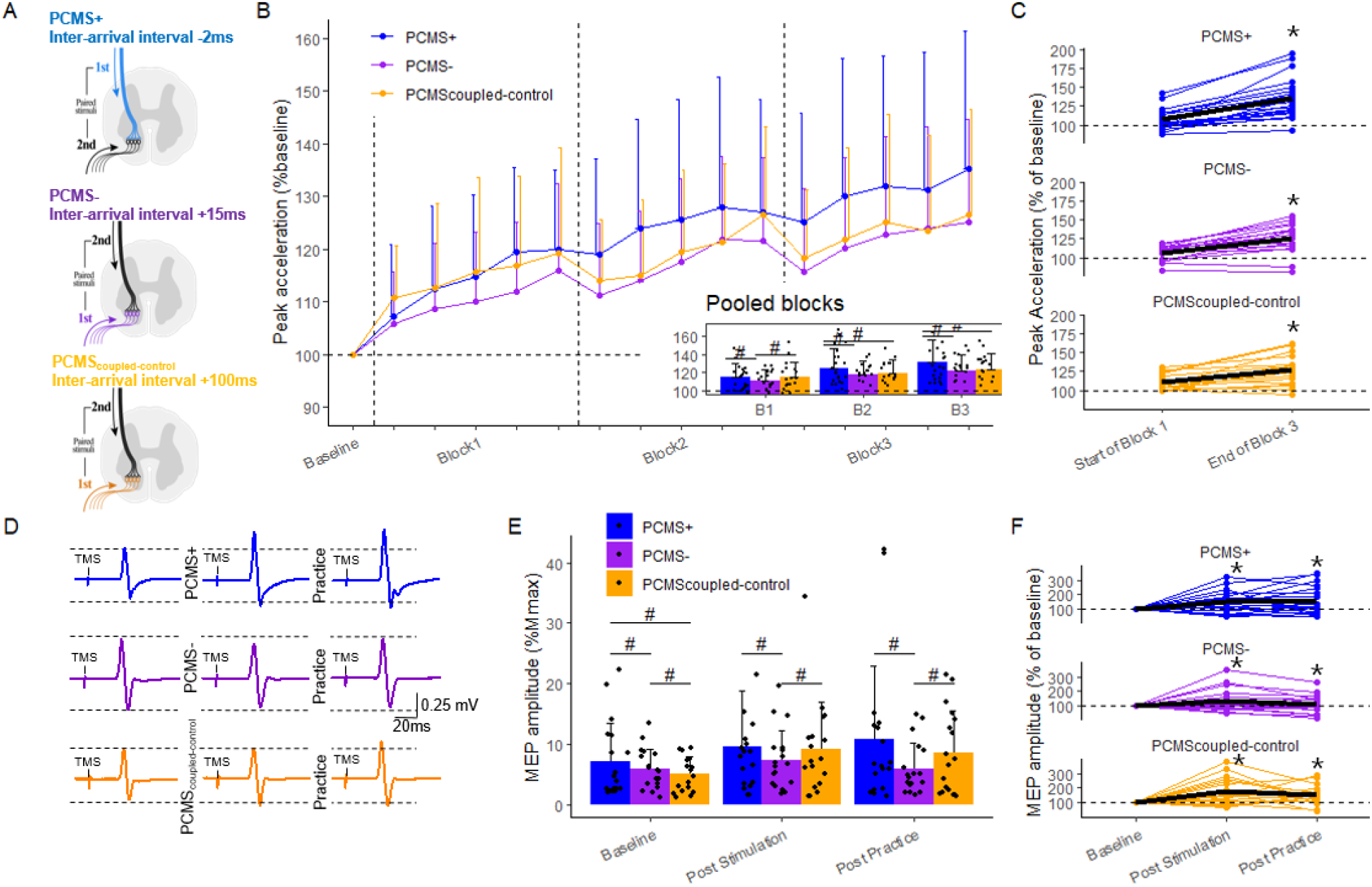
Experiment III: Effects of different PCMS protocols on ballistic motor learning and corticospinal excitability. **A)** Within-subject design, 18 participants completed three similar test days with different PCMS stimulation protocols timed according to the arrival interval at CM-synapse: −2ms (blue, PCMS+), +15 ms (purple, PCMS-), +100ms (orange, PCMS_coupled-control_), negative inter-arrival interval denotes that the TMS volley arrives at the spinal level before the PNS volley. **B)** Learning curves after the three PCMS conditions, each datapoint represent mean of 10 trials relative to individual mean at baseline. The insert shows group means across entire practice blocks with individual data depicted in dots. **C)** Individual data from the three PCMS conditions showing the mean of the 10 first trials of practice, compared to the mean of the last 10 trials of practice. The black line represents the mean performance across participants. **D)** Example of single subject data (same individual, different test days), raw MEP traces after each PCMS condition. **E)** Motor evoked potentials in percentage of M_max_ amplitude was used to quantify corticospinal excitability. **F)** Acute individual responses (MEP % baseline) to the three PCMS conditions. The black line represents the mean MEP amplitudes across participants. #p<0.05 indicates significant between-group comparison. *p<0.05 indicates significant within-group comparison.

Aligned with Experiment I and II, PCMS+ facilitated motor learning as shown from differences in performance between this and PCMS_coupled-control_ in Block 2 and 3 but not Block 1 (Figure 3B). Additionally, we found PCMS-to slow motor learning evident as a lower performance in Block 1 compared to PCMS_coupled-control_. This was supported by a LMM revealing a significant interaction effect of PROTOCOL and TIME (F_(6,8482)_=12.2, p<0.001) on peak acceleration. Comparison between protocols revealed greater improvements from B1 to B3 in ballistic performance in PCMS+ as compared to the other two protocols (PCMS+: 15.93%±0.8 vs PCMS-: 10.82%±0.8, p<0.001; and vs PCMS_coupled-control_ 8.26%±0.8, p<0.001) (Figure 3B-C). In addition, lower performance was observed during the first practice block after PCMS- as compared to the two other protocols (PCMS-: 110.5%±1.4; PCMS+: 114.4%±1.4; PCM_Scoupled-control_: 114.3%±1.4, See figure 3B).

As expected, we found facilitating effects of PCMS on corticospinal excitability as seen from increases in MEP amplitudes (Figure 3D-E). This was supported statistically by a LMM that showed a significant PROTOCOLxTIME interaction (F_(2,4143)_=10.95, p<0.001). Post-hoc comparisons revealed that PCMS+ had higher MEPs (relative to M_max_) at baseline compared to PCMS- (p<0.001) and PCMS_coupled-control_ (p<0.001). Despite of this baseline difference, PCMS+ induced relatively larger increases in MEP from baseline to post stimulation compared to PCMS- (PCMS+:51.3%±5.9; vs PCMS-: +35.3%±5.9, p=0.03), but not compared to PCMS_coupled-control_ (75.1%±6.0, p<0.01).

Experiment III confirmed the findings from the two first experiments in that facilitatory PCMS targeting the CM synapse primes subsequent ballistic motor learning and increases corticospinal excitability. The results from Experiment III additionally demonstrate that the positive effects of PCMS on motor learning are specific to protocols targeting the CM synapses inside a critical window of STDP induction and do not simply depend on arbitrary pairing of cortical and peripheral stimulation (PCMS+ vs PCMS_coupled-control_).

The results also demonstrate that PCMS effects on motor learning are bidirectional: adjusting the inter-arrival interval to one previously demonstrated to induce inhibitory after-effects on corticomotoneuronal transmission impaired early learning (PCMS-) evident from the lower performance compared to the PCMS_coupled-control_ in the first practice block. Thus, the behavioral effects conform to Hebbian learning rules in that they are contingent on temporal proximity and governed by order of spike-timing, and resemble the acute effect of PCMS previously reported for low intensity force production (Taylor & Martin, 2009). In summary, the priming effects of PCMS on ballistic motor *learning* are bidirectional depending on the spike-timing.

### Underlying neural mechanisms of PCMS and ballistic motor performance

In the present study we have replicated the line of evidence that PCMS protocols with an inter-arrival interval at the CM synapse of −2ms acutely facilitate corticospinal excitability (Bunday et al., 2018; Bunday & Perez, 2012; Christiansen et al., 2018, 2021; Jo & Perez, 2020; Taylor & Martin, 2009). Compelling evidence suggests that the increased corticospinal excitability following PCMS+ (inter-arrival interval (IAI) of 2ms) is due to enhanced transmission at the spinal level and more specifically at the corticomotoneuronal synapses. Previous studies have reported increased size of cervicomedullary motor evoked potentials (CMEPs)(Taylor & Martin, 2009). Based on the narrow spike-width in post-stimulus time histograms of single-motor unit firing after cervicomedullary electrical stimulation, the spinal motoneurons are thought to be excited predominantly through monosynaptic connections (Petersen et al., 2002). In support, PCMS has not been demonstrated to increase F-wave persistence or amplitude (indirect measures of intrinsic α-motoneuronal excitability) rendering the CM-synapse a likely site responsible for the facilitation of corticospinal excitability (Bunday & Perez, 2012). Importantly, previous work also suggests that improvements in ballistic motor performance depend on plastic changes at the corticomotoneuronal synapses evident from increases in CMEP amplitude after practice (Giesebrecht et al., 2012) along with limited changes in F-waves (Giesebrecht et al., 2011). Taken together, evidence suggests that improving ballistic practice is contingent on experience-dependent plasticity at CM synapses that are also the target of the PCMS+ protocol.

In Experiment III we compared PCMS+ to two other PCMS protocols designed to leave transmission at the CM synapse unchanged (PCMS_coupled-control_) or designed with a proposed inhibitory effect at the spinal level (PCMS-). Interestingly, we observed that PCMS_coupled-control_ led to increased MEP amplitudes, providing first evidence that repeated pairing of strong PNS and TMS at this interval increases corticospinal excitability. In conventional paired associative stimulation (PAS, Stefan et al., 2000), paired stimulations are timed with an interstimulus interval (ISI) of ~25ms to target the sensorimotor cortical circuitry mediating short latency afferent inhibition. However, it is less well studied how peripheral nerve stimulation with an ISI of ~107ms (CM IAI of 100ms) before TMS modulates the size of the MEP (Turco et al., 2018). The one study investigating this ISI found an inhibition of the MEP (Kotb et al., 2005) ascribing the effect to networks associated with long latency afferent inhibition (LAI, see e.g. Turco et al. 2018, for discussion). Although caution is warranted when concluding on the mechanisms underlying the plasticity effects of PCMS_coupled-control_ (IAI 100ms), we suggest changes in the cortical sensorimotor network mediating LAI to account for the observed facilitation. Importantly, we found that increasing corticospinal excitability by targeting the cortical level did not affect subsequent ballistic learning. Furthermore, we did not find a suppressive effect of PCMS- on corticospinal excitability. Indeed, no significant changes were observed in MEP amplitudes after PCMS-. At first glance this result is at odds with previous reports, but in contrast to Taylor & Martin (2009) we evaluated the outcome with TMS evoked MEPs and not CMEPs. Interestingly, when targeting FDI, the average interstimulus interval (ISI) needed to ensure the inter-arrival interval of 15ms at CM level was ~22.5ms, closely resembling the ISI used in facilitatory PAS, demonstrated to induce STDP-like plasticity on the cortical level (Stefan et al., 2000). It is well-known that the MEP size reflects excitability along the corticomotor signaling pathway as well as upstream of M1 (Bestmann & Krakauer, 2015). In this light, the MEP amplitude after PCMS-may reflect the sum of cortical and spinal plasticity with opposing signs. However, as we did not evaluate changes in muscle response to electrical cervicomedullary stimulation, we are limited to speculating on the loci of PCMS induced changes in excitability. Another alternative is that the PNS or TMS alone rather than the coupling *per se* caused the observed after-effects. However, as depicted in the supplementary information (Figure S2), we conducted a set of control experiments showing that neither high intensity PNS nor high intensity TMS at 0.1 Hz changed MEP amplitudes. Furthermore, we did not observe suppression of the combined MEP and F-wave during the PCMS-conditioning as compared to the PCMS_coupled-control_ (see Figure S1F, and Figure S3). The lack of suppression has previously been suggested as a marker for electrophysiological non-responders to inhibitory PCMS (Urbin et al., 2017). In summary, whereas the effects on ballistic learning displayed typical STDP-like properties, the MEP facilitation was coupling-dependent with less emphasis on the timing.

### Clinical Perspectives

Previous studies have demonstrated the clinical potential of PCMS in the recovery of dexterous hand function after spinal cord injury (Bunday & Perez, 2012) and stroke (Urbin et al., 2021). Our findings suggest that promising results of PCMS in improving dexterity in spinal cord injury may generalize to movements with other control policies through interactions between PCMS-induced and practice-dependent plasticity. The fact that the improvements persisted after practice seven days later suggests them to be therapeutically relevant. Whether repeated sessions of priming practice of brisk movements with PCMS+ result in cumulative therapeutic effects are yet to be investigated. Notwithstanding, our results support PCMS as an add-on therapy in neurorehabilitation benefitting both manipulative and vigorous movements.

## Methods

### Participants

All the experimental procedures were approved by the local ethics committee for the Greater Copenhagen area (protocol H-17019671) and the study was performed in accordance with the declaration of Helsinki. The study included fifty-eight young (28 males; age 20-30 y/o −25±4 mean ± standard deviation (SD)) adults who all volunteered and provided consent after thorough information of the study procedures. Participants were defined as able-bodied based on a standardized general eligibility questionnaire, with no history of neurological, psychiatric, or medical diseases and no intake of medication. All participants were right-handed (except one who had no preference), according to the Edinburgh Handedness Inventory (Laterality quotient: 88.21 ± 24.90 mean ± SD)(Oldfield, 1971).

### Experimental design

Three separate experiments were conducted: Experiment I, a between-group design with twenty-six participants randomized to two groups PCMS+Practice and Rest+Practice. Experiment II, a double-blinded between-group design, with twenty participants randomized to PCMS+Practice and Sham+Practice. Experiment III, a within-subject repeated measures design, where eighteen participants (incl. first 6 from PCMS group from experiment I) took part in three test days in a randomized and counterbalanced order with one week between sessions (Figure 4). A test session consisted of one of three different PCMS protocols using TMS and PNS, followed by ballistic motor training. In all experiments, participants were asked to fill in a sleep diary with their night sleep prior to the experiment. Repeated measures of motor performance (peak acceleration), corticospinal excitability (MEPs) and neuromuscular excitability (M_max_) were recorded at baseline, post-PCMS and post-ballistic training on each test day. The procedure on test days identical except for the electrophysiological intervention (i.e. paired stimulations or none) with different settings of the inter-arrival interval (IAI) in the PCMS protocols. This experimental design allowed us to investigate acute effects of different PCMS-protocols on ballistic motor performance and on the performance gains during practice of a ballistic motor task. Furthermore, it allowed us to investigate effects of different PCMS protocols on corticospinal excitability.

### Ballistic motor performance and learning

To assess ballistic motor learning, participants practiced a task requiring rapid accelerations via flexions of their right index finger. Participants were seated in a height-adjustable chair in front of a computer screen (1920×1200 res, Dell U2415) showing a white screen with a green vertical midline. Their right arm was elbow flexed approximately 90° degrees resting on the table while grabbing a custom build handle (Figure 1 A). Their index finger was placed in a metal splint placed perpendicular to the axis of rotation of the handle, with an accelerometer mounted on top of the metal splint. The handle allowed flexions of the index finger, but restricted movements in other planes. The accelerometer signal was amplified and filtered (low pass 20 Hz) and sampled at 1 kHz on a computer with a USB6008 DAQ Board (National Instruments, Inc.). A trial consisted of a rapid finger flexion within a 1 second window, with a new trial every 4^th^ second (software created for the purpose in MATLAB R2012b, MathWorks inc.). In a trial, participants were paced by a horizontal blue trace running from left to right on the white computer screen, when near the green midline the participants performed the movement. Instruction of the task consisted of a visual presentation that underlined that the movement should be performed approximately around the green midline, meaning that accuracy and precise timing at the green midline was not important. A goniometer was attached to the handle above the metacarpophalangeal joint to measure the position of the movements, and to make sure that participants got back to the stretched starting position after each trial. Participants were allowed 5 familiarization trials before the baseline test on the first test day. Motor performance was quantified as peak acceleration and was measured in 10 trials at each time point (baseline, post-PCMS and post-practice). At baseline and post-tests no augmented feedback was provided on motor performance. During practice, participants performed 3 blocks of 50 trials with a 2-minute break in between blocks. Augmented feedback was provided during the three practice blocks following each trial as a score normalized to the score measured at post-PCMS. The augmented feedback was presented as knowledge of results for 2 s before the next trial began. Additionally, verbal encouragement was provided during practice at least every 10^th^ trial to ensure that participants were motivated.

### Electrophysiological recordings

Electromyography (EMG) was recorded from m. FDI on participants’ right hand through surface electrodes (Ag-AgCl, 1 cm diameter) applied on the skin after preparation with medical sandpaper. The electrodes were placed in a muscle-belly-tendon montage, with the active electrode on the muscle belly. A zinc plate was used as ground electrode placed at the base of the hand. The EMG signals were amplified (x200), filtered (band-pass 5 Hz to 1 kHz), and sampled at 2kHz on a computer for offline analysis (Cambridge Electronic Design 1401 with Signal software v6.05). Line noise (50 Hz) was removed with a Hum Bug noise eliminator (Digitimer). During the ballistic motor task, EMG signals were amplified (x200), filtered (band-pass 5 Hz to 1 kHz), and sampled at 2kHz on a computer for offline analysis (Cambridge Electronic Design 1401 with Spike2 software v7.10).

### Transcranial magnetic stimulation

We assessed corticospinal excitability as the peak-to-peak EMG amplitudes of TMS-evoked muscle responses, called motor evoked potentials (MEPs). Monophasic single-pulse TMS was applied to the contralateral M1 to the dominant hand via a figure-of-eight TMS coil (Magstim^®^D70^2^ connected to a Magstim^200^). The hotspot for each participant in each experiment was localized via a mini-mapping procedure by determining the site of stimulation that provided large and robust MEP responses in the FDI. The coil was placed with the centre oriented parallel to the scalp over the hotspot of FDI representation with the handle of the coil pointing backward at an angle of 45° to the sagittal and horizontal axis (Figure 5A). This induces a posterior-anterior current direction in the targeted cortex. The resting motor threshold (rMT) was defined as the stimulus intensity needed to elicit recognizable MEPs with an amplitude above 0.05 mV in 5/10 consecutive stimulations (Rossini et al., 2015). Muscle relaxation was monitored through concurrent EMG recordings. Twenty TMS stimulations (120% of rMT) were delivered at each time point (2xbaseline, post-PCMS and post motor practice) to measure MEP amplitudes. A neuro-navigation system (Brainsight 2, Rogue Research, Montreal, Canada) was used to ensure stable positioning of the coil throughout the experiments. MEP latencies during voluntary contraction (10%MVC) were recorded to calculate the central conduction time used to inform the interstimulus intervals in the paired stimulations protocols.

### Electrical peripheral nerve stimulation

Electrical stimulation with high voltage electrical current (200 μs pulse duration, DS7A; Digitimer) was delivered to the ulnar nerve at the wrist (Bar Stimulating Electrode, Digitimer) to measure the maximal compound muscle action potential (M_max_) and F-waves. The latencies of M_max_ and F-waves were used to calculate the peripheral conduction time used to individualize the paired stimulations procotols (Figure 4S, B-D). F-wave latency was defined as the F-wave with the earliest onset (Christiansen et al., 2018).

### Paired corticomotoneuronal stimulation protocols

PCMS protocols were individualized based on recordings of M_max_, Fwave and MEP latencies from each participant on each test day. Interstimulus intervals were based on calculations of individual peripheral and central conduction times. The stimulation protocols targeted the corticospinal-motoneuronal synapses with different inter-arrival intervals between the descending TMS volley and the antidromic stimulation from PNS (see supplementary information, Figure S1 for details).

### Quantification and Statistical analysis

Visual inspection of all signals during and after the experiments, ensured exclusion of M-waves and MEPs that contained pre-stimulus EMG activity 100 ms before stimulation (see Rogasch et al., 2009). Post hoc analysis included removal of outliers defined as mean±2 standard deviation for the given measurement. MEPs were normalized to the respective M_max_ amplitude to allow between-session comparisons. Peak acceleration scores were normalized to the baseline value obtained in the same experimental session to allow comparisons between participants. Data from the motor practice sessions a total of 150 trials of peak acceleration was binned in trials of 10 to analyze motor performance within a practice session.

All statistical analyses were performed using R (R Core Team, 2022, version 4.1.3). Linear mixed effect models were fitted to data for all dependent variables (peak acceleration, MEP amplitudes) using the *lme4* R-package (Bates et al., 2014). For Experiment I and II, linear mixed effect models with the fixed factors GROUP (2 levels: Rest/Sham and PCMS) and TIME (Experiment I: 4 levels: Baseline, Block1-3, Experiment II: 5 levels: Baseline, Block1-3, Day7) was fitted to the acceleration data with an interaction term (GROUP x TIME). In Experiment III, linear mixed effect models with the fixed factors PROTOCOL (3 levels: PCMS+; PCMS-; PCMS_coupled-control_) and TIME (4 levels: Baseline, Block1-3) were fitted to the acceleration data with an interaction term (GROUP x TIME). Potential order effects were accommodated statistically by adding ‘DAY’ to the model with an additive term. Random intercepts were fitted for each subject to account for the repeated measures design. Assumptions of normality and homogeneity of variance of residuals were inspected by quantile-quantile plots and residual plots. To evaluate significance of main effects or interactions, we used the R-package *lmerTest* (A, Kuznetsova, et al. 2017) that computes P-values from mixed effect models via the Satterthwaite’s degrees of freedom method. If main effects or interactions were significant, we proceeded to pairwise comparisons using the *multcomp* R-package (Hothorn et al., 2008). Namely, we computed contrasts to test specific hypotheses, e.g., how ballistic performance was affected after the paired stimulations (during practice). These contrasts are presented as model estimates with standard errors (SE). For all post-hoc comparisons, the Holm-Sidak method was used to adjust for multiple statistical comparisons. For all statistical analyses the significance level was set at p<0.05.

## Acknowledgements

The authors would like to thank the participants for their time and patience. We are grateful for the financial support from Nordea-fonden (Grant no. 02-2019-00045). Lasse Christiansen holds a postdoc grant from the Lundbeck Foundation (Grant no. R322-2019-2406). Mikkel Malling Beck holds a postdoc grant from the Capital Region of Copenhagen (Region H) and was funded by a grant from Innovation Fund Denmark (Innovation Fund Denmark, grant no. 9068-00025B) and the Danish Ministry of Culture (Grant no. FKP.2018-0070).

## Author contributions

Conceptualization, J.L.J., M.M.B., L.C., and J.R.B.; Methodology, J.L.J., M.M.B. and J.R.B.; Formal Analysis, J.R.B. and M.M.B.; Investigation, J.R.B., L.J. and M.M.B.; Writing – Original Draft, J.R.B.; Writing – Review & Editing, J.R.B., M.M.B., L.C., L.J. and J.L.J.; Visualization, J.R.B.; Supervision, M.M.B. and J.L.J. Project Administration, J.L.J.; Funding Acquisition, J.L.J.

## References

Bestmann, S. & J.W. Krakauer (2015): The uses and interpretations of the motor-evoked potential for understanding behaviour. Experimental Brain Research, Vol. 233:3, pp. 679–689.

Bi, G.Q. & M.M. Poo (1998): Synaptic modifications in cultured hippocampal neurons: Dependence on spike timing, synaptic strength, and postsynaptic cell type. Journal of Neuroscience, Vol. 18:24, pp. 10464–10472.

Bunday, K.L. & M.A. Perez (2012): Motor recovery after spinal cord injury enhanced by strengthening corticospinal synaptic transmission. Current Biology, Vol. 22:24, pp. 2355–2361.

Bunday, K.L., M.A. Urbin & M.A. Perez (2018): Potentiating paired corticospinal-motoneuronal plasticity after spinal cord injury. Brain Stimulation, Elsevier Ltd, Vol. 11:5, pp. 1083–1092.

Christiansen, L., B. Chen, Y. Lei, M.A. Urbin, M.S.A. Richardson et al. (2021): Acute intermittent hypoxia boosts spinal plasticity in humans with tetraplegia. Experimental Neurology, Elsevier, Vol. 335:May 2020, p. 113483.

Christiansen, L. & M.A. Perez (2018): Targeted-Plasticity in the Corticospinal Tract After Human Spinal Cord Injury. Neurotherapeutics, Neurotherapeutics, Vol. 15:3, pp. 618–627.

Christiansen, L., M.A. Urbin, G.S. Mitchell & M.A. Perez (2018): Acute intermittent hypoxia enhances corticospinal synaptic plasticity in humans. ELife, Vol. 7:, pp. 1–17.

D’Amico, J.M., S.C. Dongés & J.L. Taylor (2018): Paired corticospinal-motoneuronal stimulation increases maximal voluntary activation of human adductor pollicis. Journal of Neurophysiology, Vol. 119:, pp. 369–376.

Dongés, S.C., C.L. Boswell-Ruys, J.E. Butler & J.L. Taylor (2019): The effect of paired corticospinal–motoneuronal stimulation on maximal voluntary elbow flexion in cervical spinal cord injury: an experimental study. Spinal Cord, Springer US, Vol. 57:9, pp. 796–804.

Giesebrecht, S., H. Van Duinen, G. Todd, S.C. Gandevia & J.L. Taylor (2012): Training in a ballistic task but not a visuomotor task increases responses to stimulation of human corticospinal axons. Journal of Neurophysiology, Vol. 107:, pp. 2485–2492.

Giesebrecht, S., P.G. Martin, S.C. Gandevia & J.L. Taylor (2011): Altered corticospinal transmission to the hand after maximum voluntary efforts. Muscle and Nerve, Vol. 43:5, pp. 679–687.

Jo, H.J. & M.A. Perez (2020): Corticospinal-motor neuronal plasticity promotes exercise-mediated recovery in humans with spinal cord injury. Brain, pp. 1–15.

Kotb, M.A., T. Mima, Y. Ueki, T. Begum, A.T. Khafagi et al. (2005): Effect of spatial attention on human sensorimotor integration studied by transcranial magnetic stimulation. Clinical Neurophysiology, Vol. 116:5, pp. 1195–1200.

Kuznetsova, A., P.B. Brockhoff, R.H.B. Christensen & S.P. Jensen (2017): lmerTest package: Tests in Linear Mixed Effects Models. Journal of Statistical Software, Vol. 82:13, pp. 1–26.

Oldfield, R.C. (1971): The assessment and analysis of handedness: The Edinburgh inventory. Neuropsychologia, Vol. 9:, pp. 97–113.

Petersen, N.T., J.L. Taylor & S.C. Gandevia (2002): The effect of electrical stimulation of the corticospinal tract on motor units of the human biceps brachii. Journal of Physiology, Vol. 544:1, pp. 277–284.

Rogasch, N.C., T.J. Dartnall, J. Cirillo, M.A. Nordstrom & J.G. Semmler (2009): Corticomotor plasticity and learning of a ballistic thumb training task are diminished in older adults. Journal of Applied Physiology, Vol. 107:6, pp. 1874–1883.

Rossini, P.M., D. Burke, R. Chen, L.G. Cohen, Z. Daskalakis et al. (2015): Non-invasive electrical and magnetic stimulation of the brain, spinal cord, roots and peripheral nerves: Basic principles and procedures for routine clinical and research application: An updated report from an I.F.C.N. Committee. Clinical Neurophysiology, Vol. 126:6, pp. 1071–1107.

Shmuelof, L. & J.W. Krakauer (2011): Are we ready for a natural history of motor learning? Neuron, Elsevier Inc., Vol. 72:3, pp. 469–476.

Shulga, A., P. Lioumis, E. Kirveskari, S. Savolainen, J.P. Mäkelä et al. (2015): The use of F-response in defining interstimulus intervals appropriate for LTP-like plasticity induction in lower limb spinal paired associative stimulation. Journal of Neuroscience Methods, Elsevier B.V., Vol. 242:, pp. 112–117.

Siebner, H.R., K. Funke, A.S. Aberra, A. Antal, R. Chen et al. (2022): Transcranial magnetic stimulation of the brain: What is stimulated?-a consensus and critical position paper. Clinical Neurophysiology, International Federation of Clinical Neurophysiology, available at:http://doi.org/10.1016/j.clinph.2022.04.022.

Sinopoulou, E., E.S. Rosenzweig, J.M. Conner, D. Gibbs, C.A. Weinholtz et al. (2022): Rhesus macaque versus rat divergence in the corticospinal projectome. Neuron, The Authors, Vol. 110:18, pp. 2970–2983.e4.

Stefan, K., E. Kunesch, L.G. Cohen, R. Benecke & J. Classen (2000): Induction of plasticity in the human motor cortex by paired associative stimulation. Brain, Vol. 123:3, pp. 572–584.

Taylor, J.L. & P.G. Martin (2009): Voluntary motor output is altered by spike-timing-dependent changes in the human corticospinal pathway. Journal of Neuroscience, Soc Neuroscience, Vol. 29:37, pp. 11708–11716.

Team, R.C. (2022): R: A language and environment for statistical computing, R Foundation for Statistical Computing, Vienna, Austria., available at: https://www.r-project.org/.

Turco, C. V., J. El-Sayes, M.J. Savoie, H.J. Fassett, M.B. Locke et al. (2018): Short-and long-latency afferent inhibition; uses, mechanisms and influencing factors. Brain Stimulation, Elsevier Ltd, Vol. 11:1, pp. 59–74.

Urbin, M.A., J.L. Collinger & G.F. Wittenberg (2021): Corticospinal recruitment of spinal motor neurons in human stroke survivors. Journal of Physiology, Vol. 599:18, pp. 4357–4373.

Urbin, M.A., R.A. Ozdemir, T. Tazoe & M.A. Perez (2017): Spike-timing-dependent plasticity in lower-limb motoneurons after human spinal cord injury. Journal of Neurophysiology, Vol. 118:4, pp. 2171–2180.

