## Supplementary Information for "Hebbian priming of human spinal motor learning"

### Paired corticomotoneuronal stimulation protocols

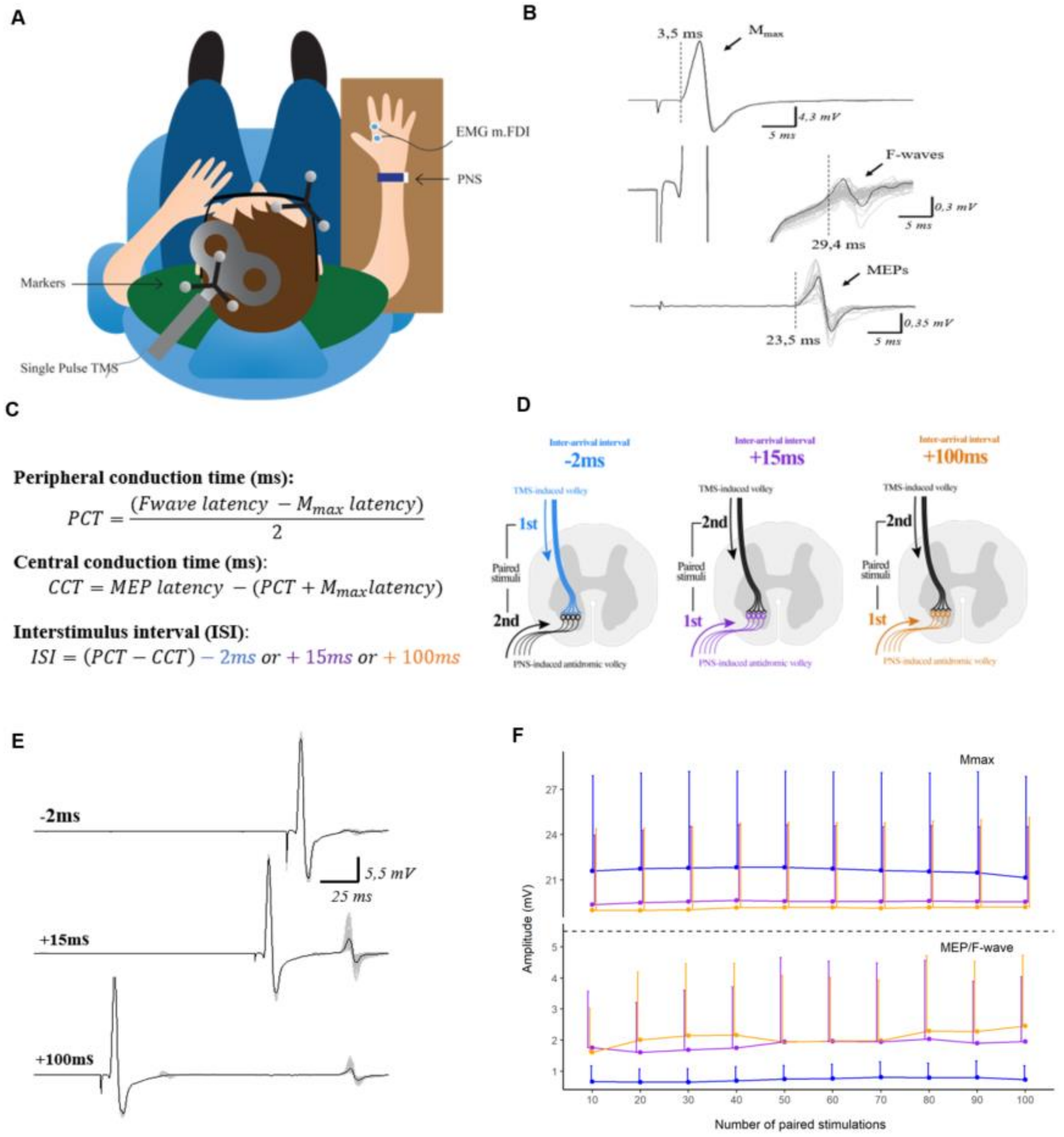


**Figure S1. Plasticity-inducing protocols**. **A:** Participants at rest while receiving the paired stimulations from transcranial magnetic stimulation (TMS) and electrical peripheral nerve stimulation (PNS), markers on TMS coil and participants were calibrated with Brainsight Software. **B:** Individual latencies from one participant: raw traces of $M_{max}$, Fwaves and MEP were obtained on each test day for each participant. **C:** Interstimulus intervals were based on calculations of individual peripheral and central conduction times. **D:** The stimulation protocols targeted the corticospinal-motoneuronal synapses with different inter-arrival intervals between the descending TMS volley and the antidromic stimulation from PNS. **E:** Visualizing the different intervals with overdraw traces (100 paired stimulations) from each PCMS protocol (-2ms, +15ms and +100ms) from a single participant. **F:** Graph shows the pooled data of $M_{max}$ and MEPs during the PCMS protocols; -2ms (blue), +15ms (purple), and +100ms (orange). Each data point represents the average of 10 traces ($M_{max}$ and MEP's). Note the separation of the y-axis. Error bars indicate SD.

### Effects of non-paired rPNS or rTMS on corticospinal excitability

Ten participants from Experiment III also completed two extra control sessions. Since all the PCMS protocols in Experiment III led to increase in MEP response, we conducted a control experiment, to investigate whether non-paired low frequency TMS or PNS alone could be responsible for the observed increase in corticospinal excitability. This was not the case since neither of the non-paired low frequency (0.1Hz) protocols led to increase in MEPs as measured following stimulation of the M1 with an intensity corresponding to 120% RMT (Figure S2). This was supported statistically with no change in normalized MEP amplitudes across 45 minutes after both rPNS (one-way ANOVA, $F_{(4,1030)}=1.804$, $p=0.13$) and rTMS (one-way ANOVA, $F_{(4,970)}=0.751$, $p=0.55$).

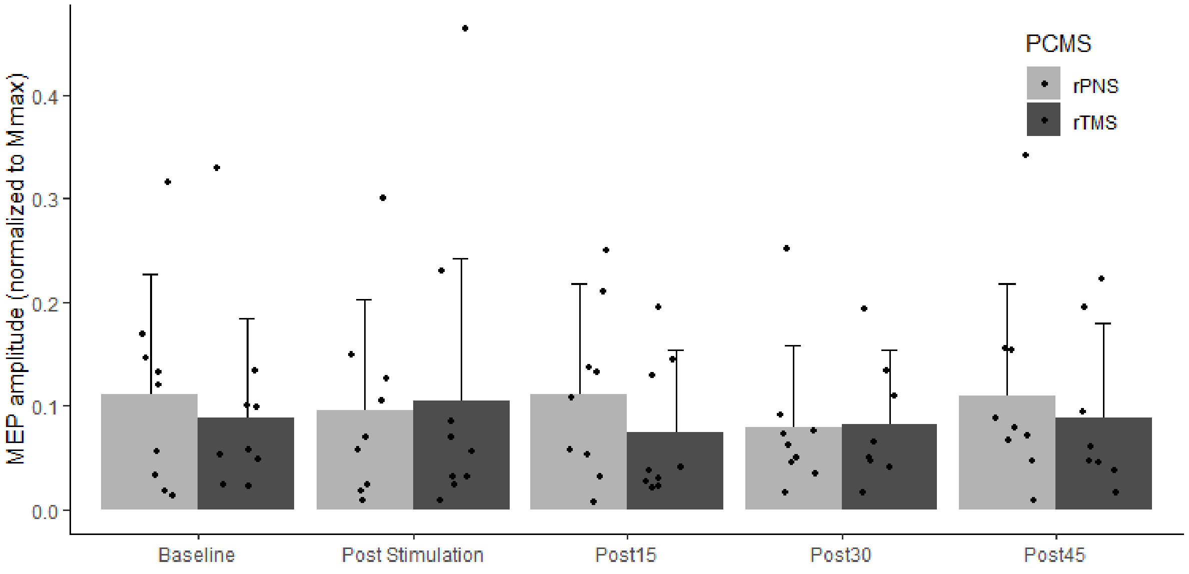


**Figure S2. Effects of non-paired low frequency stimulations;** MEP amplitudes normalized to Mmax amplitude after rPNS (lightgrey) and rTMS (darkgrey). Barchart represent group means+SD and each datapoint is individual means at given timepoint.

### Responders/Non-Responders in PCMS-

Only 6 out of 18 participants showed decreased MEP amplitude after the PCMS-protocol. MEP amplitudes during PCMS- have previously been proposed as a marker for electrophysiological responders and non-responders to inhibitory PCMS- (Urbin et al., 2017). Participants were binary categorized as responders to PCMS- if MEP amplitude was decreased post PCMS-. Visually, data shows that participants with decreased MEPs after PCMS- (responders) displayed smaller MEP amplitudes during the PCMS- protocol (even relatively to baseline that was acquired at a lower stimulation intensity than used during PCMS: 120% RMT vs. 150% RMT, respectively) (Figure S3). It could further be speculated that MEP amplitudes during PCMS- in those responding with increases in MEP amplitudes following PCMS- (non-responders) were close to MEPmax (this is likely, based on the TMS intensity during PCMS, 150% rMT), and thus in line with Urbin et al. (2017).

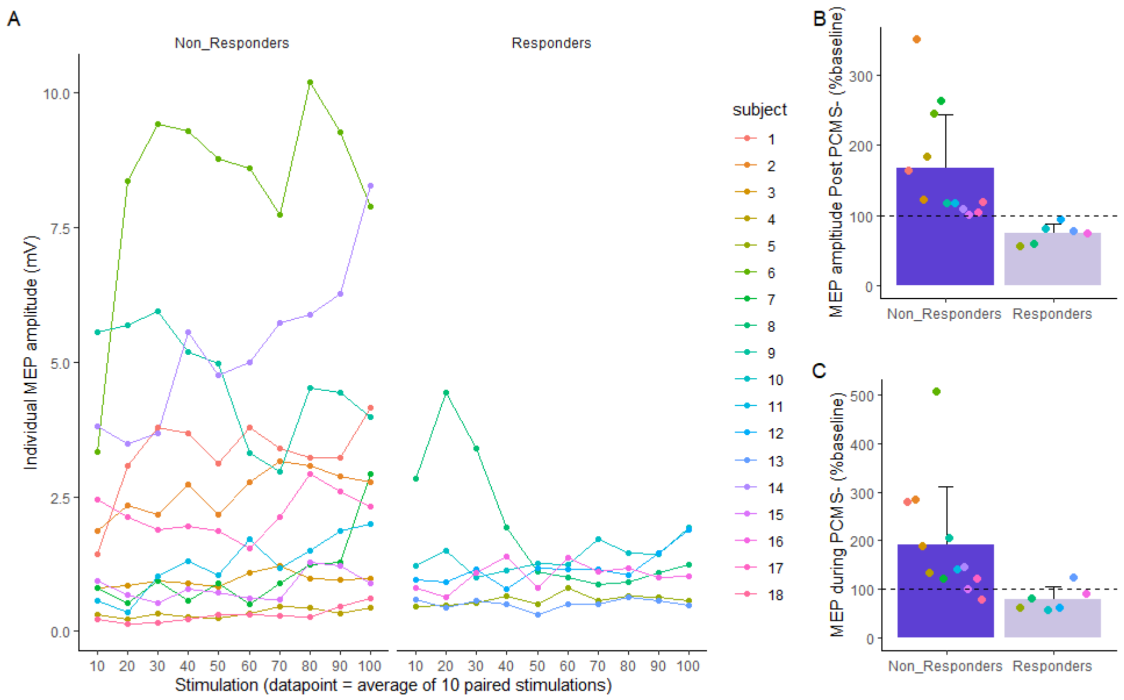


**Figure S3. Responders vs Non-responders to PCMS-. A)** MEP amplitudes during PCMS-, divided in whether participants responded as intended to the PCMS- (decreased MEP amplitude post PCMS-). Each datapoint is the individual mean of 10 stimulations. **B)** Group and individual means of MEP amplitudes during all stimulations delivered during PCMS- presented as % of MEP amplitudes at baseline. Please note that for the majority of individuals responding as intended to the PCMS-, the MEP amplitudes delivered during PCMS- (150% RMT) was smaller than MEP amplitudes delivered at baseline (120% RMT) showing an effective reduction of MEP amplitudes.
